# The likelihood of heterogeneity or additional mutation in KRAS or associated oncogenes to compromise targeting of oncogenic KRAS G12C

**DOI:** 10.1101/149724

**Authors:** Vincent L. Cannataro, Stephen G. Gaffney, Carly Stender, Zi-Ming Zhao, Mark Philips, Andrew E. Greenstein, Jeffrey P. Townsend

## Abstract

Activating mutations in RAS genes are associated with approximately 20% of all human cancers. New targeted therapies show preclinical promise in inhibiting the KRAS G12C variant, however, concerns exist regarding the effectiveness of such therapies *in vivo* given the possibilities of existing intratumor heterogeneity or *de novo* mutation leading to treatment resistance. We performed deep sequencing of 27 KRAS G12 positive lung tumors and found no evidence of other oncogenic mutations within KRAS or within commonly mutated downstream genes that could confer resistance at the time of treatment. Furthermore, we estimate the *de novo* mutation rate in KRAS position 12 and in genes downstream of *KRAS.* We find that mutations that confer resistance are about as likely to occur downstream of KRAS as within KRAS. Moreover, we present an approach for estimation of the selection intensity for these point mutations that explains their high prevalence in tumors. Our approach predicts that BRAF V600E would provide the highest fitness advantage for *de novo* resistant subclones. Overall, our findings suggest that resistance to targeted therapy of KRAS G12C positive tumors is unlikely to be present at the time of treatment and, among the *de novo* mutations likely to confer resistance, mutations in BRAF, a gene with targeted inhibitors presently available, result in subclones with the highest fitness advantage.

**One Sentence Summary:** Mutations conferring resistance to KRAS G12C targeted therapy are unlikely to be present at the time of resection, and the likely mechanisms of evolved resistance are predicted be ones that are responsive to therapies that are in development or that are already available.

## Introduction

In roughly 20% of all human tumors, mutational activation within the *RAS* gene family is implicated. Mutations in the *KRAS* isoform are a major driver in 90% of pancreatic ductal adenocarcinoma, 43% of colorectal cancer, and 26% of non-small-cell lung cancer (Downward and Julian 2003; Eser et al. 2014; Fernandez-Medarde and Santos 2011). *KRAS* was one of the first proto-oncogenes identified (Cox and Der 2010; Chang et al. 1982) and has long been the target of drug development efforts (Cox, Der, and Philips 2015; McCormick 2015). *KRAS* gene products control the pathways involved in cell proliferation, differentiation, and apoptosis by cycling between GTP-bound active and GDP-bound inactive signaling states (Fernandez-Medarde and Santos 2011; Shima et al. 2013). Most oncogenic mutations in the *KRAS* gene change the 12th amino acid of the KRAS protein from a glycine to some other amino acid, compromising GTPase activity and resulting in persistent stimulation of mitogenic signaling (Prior, Lewis, and Mattos 2012). Design of therapeutics toward the G12 mutation has proven extremely challenging, largely because of the protein’s high affinity for GTP and insufficient knowledge of allosteric regulatory sites (Ostrem et al. 2013; Shima et al. 2013; Greulich 2010; Singh, Longo, and Chabner 2015).

Activating point mutations in *KRAS* account for a significant portion of all *RAS* mutations found in human cancers (Downward and Julian 2003) with 80% of activating mutations occurring at codon 12 (Prior, Lewis, and Mattos 2012), and with 3%, 8%, and 49% of codon position 12 substitutions resulting in a cysteine in pancreatic (PAAD), colorectal (COADREAD), and lung adenocarcinomas (LUAD), respectively (Porta et al. 2009). Patients that underwent surgical resection of a G12C positive LUAD tumor had shorter disease free survival than patients who had resections of LUAD tumors with other G12 mutations (Nadal et al. 2014) or wild-type KRAS (Izar et al. 2014; Nadal et al. 2014). Additionally, in (Downward and Julian 2003; Eser et al. 2014; Fernandez-Medarde and Santos 2011), G12C mutations are a strong negative predictor of EGFR tyrosine-kinase inhibitor therapy efficacy (Fiala et al. 2013) and is associated with shorter overall survival in second- and third-line chemotherapies (Svaton et al. 2016).

Perhaps the most promising approach targets just those mutations of the G12 codon that lead to the incorporation of a solvent-accessible cysteine residue (G12C mutations) in a novel pocket adjacent to the nucleotide binding site (Hunter et al. 2014; Patricelli et al. 2016; Hobbs et al. 2016; Visscher, Arkin, and Dansen 2016; Wilson and Tolias 2016). Design of a small molecule that can form a covalent sulfur bond at this key site would overcome the protein’s native nucleotide preference for GTP, thus eliminating the constitutive activation of the oncoprotein (Ostrem et al. 2013; Westover, Jänne, and Gray 2016; Hunter et al. 2014; Montalvo, Li, and Westover 2017; Xiong et al. 2017). A major concern regarding the prospects for therapeutic success of targeted therapies in general (McGranahan and Swanton 2015; Burrell and Swanton 2014)—and of this approach specifically (Al-Mulla et al. 1998; Alsdorf et al. 2013)—has been tumor heterogeneity. If low-frequency subclones are present in KRAS G12C tumors that feature G12 mutations encoding amino acids other than cysteine—or if such mutations occurred rapidly during the course of therapy—then resistance to a sulfur-bonding G12C therapeutic would rapidly evolve in the patient, markedly diminishing and eventually eliminating the utility of the therapeutic approach.

Although both the nature and prevalence of the KRAS G12C mutation make it a promising target for covalent inhibitors, intratumor heterogeneity could limit the success of such therapeutics. Somatic aberrations within a tumor could be subject to unique microenvironmental selection pressures, leading to spatial heterogeneity of the genetics of subclones (McGranahan and Swanton 2015; Burrell et al. 2013). If a tumor possessing drug resistant subclones is subject to the selection pressures of a targeted therapy, those subclonal cells will repopulate the tumor and ultimately render therapy ineffective in cases of recurrence (Keats et al. 2012; Burrell and Swanton 2014; Merlo et al. 2006; Schmitt, Loeb, and Salk 2016). This challenge is exemplified in EGFR-driven NSCLC, in which subclones with the EGFR T790M mutation—known to decrease the effect of tyrosine kinase inhibitors (TKIs)—have been identified in up to 79% of treatment-naive tumors (Yun et al. 2008; Fujita et al. 2012; Yu et al. 2013). Pressure from TKIs select for enriched growth of T790M mutated subclones and lead to overall acquired TKI resistance (Stewart et al. 2015). Therefore, the efficacy of targeted therapy can be hindered by intra-tumor heterogeneity of the target gene present in the tumor before treatment. Indeed, a previous tumor sequencing study found that one out of 90 pancreatic ductal adenocarcinoma tumors positive for KRAS amino acid 12 mutations were heterogeneous for KRAS G12, containing both the nucleotide 34 G→T mutation responsible for the G12C mutation and the dinucleotide 34–35 GG→TT mutation resulting in G12F, in different locations of the same tumor (Hashimoto et al. 2016).

The efficacy of targeted therapy can also be hindered by drug-induced selection operating on mutations that occur after the start of treatment. *De novo* mutations can alter the target gene, preventing the binding and/or therapeutic effects of the treatment, or they can occur within genes in the downstream pathway of the target, possibly reactivating the oncogenic pathway. For instance, variant substitutions in *BRAF* have been observed in 1% of lung adenocarcinomas with acquired resistance to EGFR inhibitors, with no variant substitutions in other genes downstream of *EGFR* (Ohashi et al. 2012), and 60% of colorectal cancer patients with acquired resistance to *EGFR* inhibitors harbor variant substitutions in *KRAS* in metastases that were not detected prior to treatment (Misale et al. 2012). Although mutations at these sites have been reported to arise *de novo*, recent work suggests that somatic variants at these sites might be present as low-frequency subclones in metastatic colorectal cancer (Tougeron et al. 2013). Measuring the frequency of treatment-resistant subclones, and the likelihood that subclones may arise after treatment, is essential to predicting treatment efficacy (Ziogas et al. 2016).

Here we perform tumor sequence analysis, patient-derived xenograft (PDX) tumor evolution experimentation, and evolutionary inference to understand the potential for evolution of resistance to therapies directed toward KRAS G12C. To ascertain the potential for tumor heterogeneity to harbor alternate oncogenic KRAS G12C mutations, we performed extremely deep sequencing of 27 KRAS G12 mutant primary lung tumors, 16 of which exhibited a variant amino acid substitution at position 12 to cysteine. Using this deep sequence data, we evaluated whether heterogeneity would be likely to be present at the time of therapeutic intervention. Furthermore, we performed sequencing of common oncogenic *KRAS* mutations in two PDXs of KRAS G12C mutant tumors, passaged eight times each, to ascertain the frequency with which alternate mutations at the G12 site might arise *de novo*. Then, by calculating the mutation rates and selection coefficients for mutations within KRAS and mutations within other genes downstream of KRAS in the RAS pathway, we quantified the likelihood of other mutations that might confer resistance to treatment subsequent to commencement of therapy. Together, these analyses provide insight into the likelihood of resistance evolution in patients treated with a specific inhibitor of the G12C oncogenic function.

## Materials and Methods

To ascertain the likelihood that tumors harbor multiple mutated *KRAS* alleles, 27 human lung adenocarcinoma tumors were collected by surgical resection (Asterand Bioscience, Detroit, MI). As non-neoplastic controls, cadaveric lung tissue was obtained from three trauma victims aged 19–30 (Asterand Bioscience, Detroit, MI). Twenty-one of the tumors and all three controls were flash frozen, while six of the tumors were formalin fixed and paraffin embedded. Sequence enrichment and library preparation were performed in duplicate using SuraSeq^®^ 500 reagents, followed by sequencing on the Illumina MiSeq. DNA quality was assessed by A260/280/230 and quantitative functional index (QFI), a measure of the amplicons that are of sufficient length for next generation sequencing (NGS) analysis. NGS data quality was assessed by mate concordance and consistency across repeat MiSeq runs. The mean depth of coverage for the *KRAS* gene was 48,163× (23,922×–251,359×). In addition to the KRAS allele, a select number of regions of genes commonly mutated in cancer (Table 1) were amplified and sequenced in parallel to gauge the overall frequency and heterogeneity of relevant mutations in each tumor.

**Table 1:**
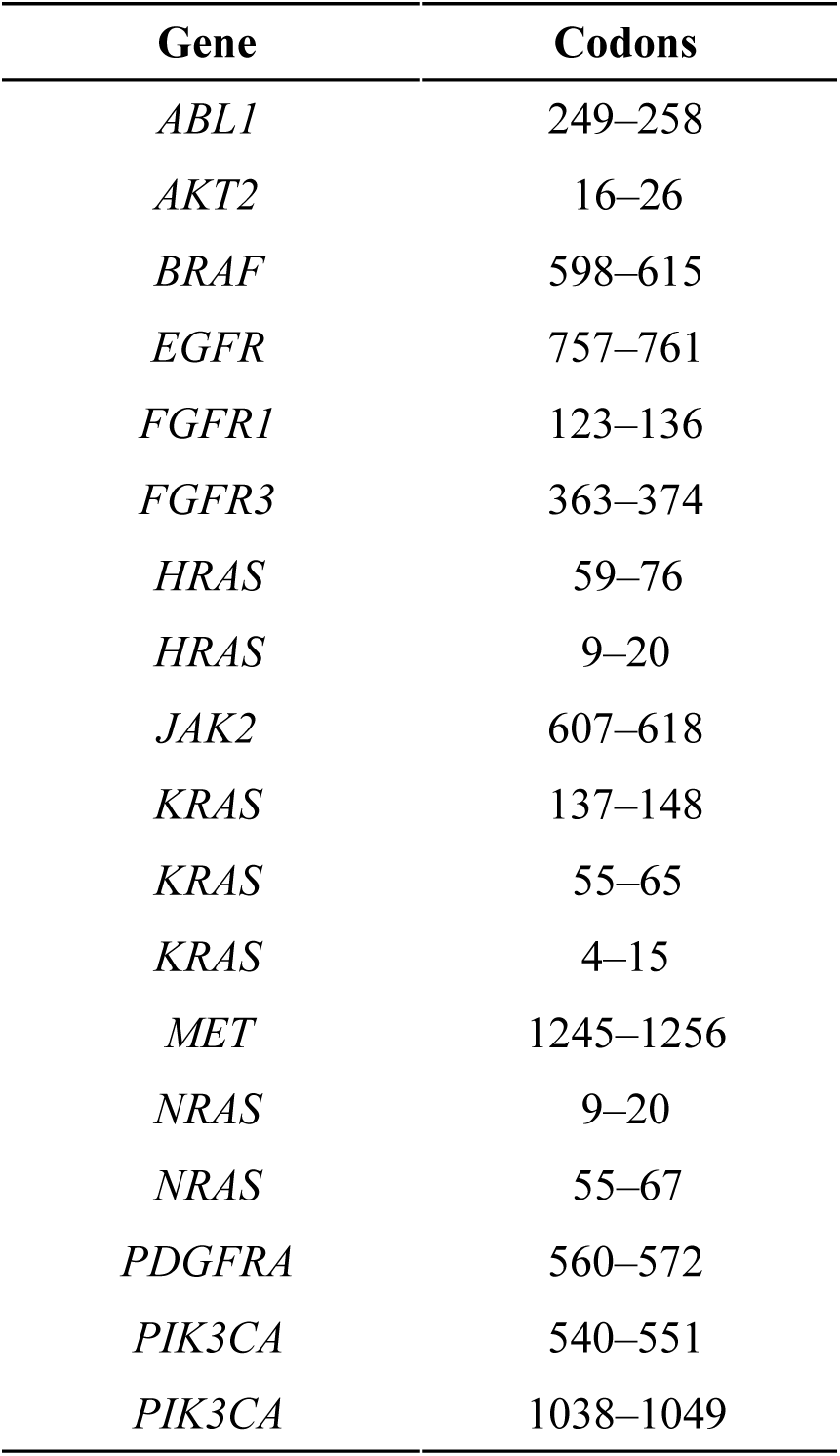
Gene locations sequenced.

| **Gene** | **Codons** |
| --- | --- |
| *ABL1* | 249–258 |
| *AKT2* | 16–26 |
| *BRAF* | 598–615 |
| *EGFR* | 757–761 |
| *FGFR1* | 123–136 |
| *FGFR3* | 363–374 |
| *HRAS* | 59–76 |
| *HRAS* | 9–20 |
| *JAK2* | 607–618 |
| *KRAS* | 137–148 |
| *KRAS* | 55–65 |
| *KRAS* | 4–15 |
| *MET* | 1245–1256 |
| *NRAS* | 9–20 |
| *NRAS* | 55–67 |
| *PDGFRA* | 560–572 |
| *PIK3CA* | 540–551 |
| *PIK3CA* | 1038–1049 |

We also checked for heterogeneity within *KRAS* and within other gene sequences that are known to be focal for RAS pathway oncogenic mutations (Table 1) in whole-exome tumor-normal sequencing data curated by The Cancer Genome Atlas (TCGA) and gathered at Yale University, consisting of data from 650 lung adenocarcinoma (542 TCGA and 108 Yale–Gilead), 235 pancreatic adenocarcinoma (184 TCGA and 51 Yale–Gilead), and 489 (all from TCGA) colorectal adenocarcinoma tumors. All TCGA data was downloaded from the Broad Institute TCGA Genome Data Analysis Center (2016)(Broad Institute 2016).

To determine whether a given mutation was distinguishable from the technical noise inherent to deep sequencing experiments, we estimated the noise across the samples analyzed. We estimated noise by averaging the rate of detection of all possible mutations at *KRAS* codons 7, 8, 9, 10, 11, 14, and 15, where somatic variant substitutions are not generally observed in cancer tissues. Three standard deviations above the mean (~0.3% of reads) was used as a threshold for distinguishing a true mutation from noise. In one case, the sequencing depth was increased to 429,038×. This ultra-deep sequencing did not affect this threshold, indicating that the noise more likely originated from the DNA amplification than the sequencing reaction.

To assess the potential for novel KRAS mutations to arise on a KRAS G12C positive tumor during tumor growth, two KRAS G12C human lung adenocarcinoma PDXs were implanted subcutaneously in female immune-compromised *nu/nu* mice (Harlan) for 8 passages (Champions Oncology). Tumors were passaged after ~5 doublings in volume; PDXs were analyzed by allele-specific PCR (Diacarta Inc, Richmond, CA) at passages 2–8 (Fig. 1).

**Figure 1.**
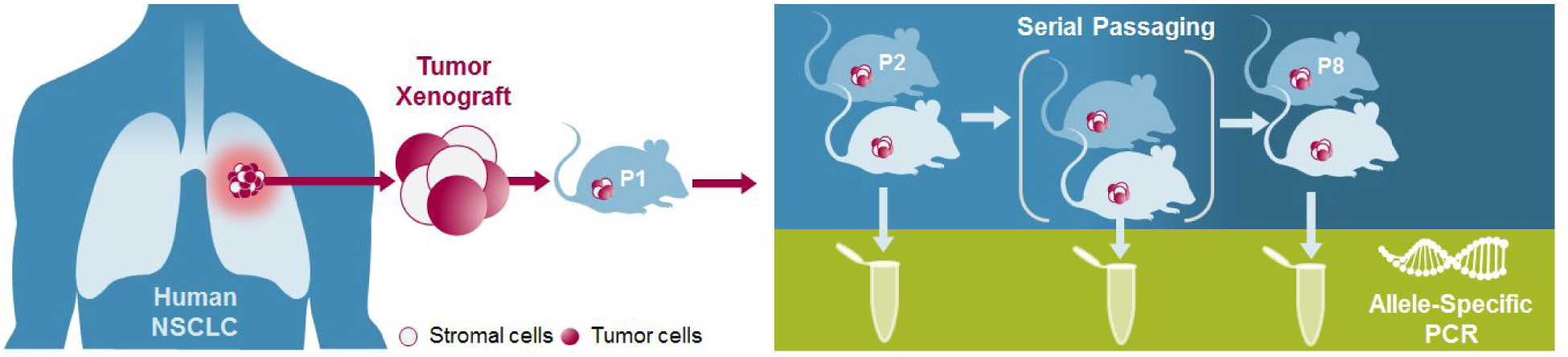
Schematic of KRAS mutation assessment in human tumor samples serially passaged in mice. The tumor and human stromal cells are resected and implanted in the flank of a BALB/c nude mouse. The tumor expands in the mouse flank and the human stromal cells are rapidly replaced with mouse stromal cells. Tumors (and contaminating stroma) are sampled after each passage.

Next, we estimated the rate at which *de novo* mutations that might compromise treatment directed at the G12C allele would arise within KRAS during treatment. In particular, we estimate the rate of new oncogenic mutations at somatic variant position 12 because they would prevent a G12C-specific inhibitor from binding, and also the rate of new oncogenic mutations at position 61 because cells with KRAS^G12C/Q61L^ have been shown to be less sensitive to the G12C-specific inhibitor ARS-853 when compared to KRAS^G12C^ mutant cells (Lito et al. 2016). The rate that single nucleotides are mutated in *KRAS* was estimated using `MutSigCV`, a software package that combines mutation data with information on chromatin state, replication timing, and transcriptional activity to calculate gene-specific mutation rates (Lawrence et al. 2013). The `MutSigCV` analysis was informed by averaged gene expression levels in lung cell lines, obtained from The Cancer Cell Line Encyclopedia (Barretina et al. 2012). The mutation dataset consisted of single nucleotide variants observed in 108 lung adenocarcinoma tumor-normal whole exome sequences gathered at Yale University and 542 publicly available lung adenocarcinoma tumor-normal whole exome sequences generated by The Cancer Genome Atlas (TCGA) Research Network (Broad Institute TCGA Genome Data Analysis Center, 2016). We evaluated the relative rates of specific nucleotide transitions and transversions based on their trinucleotide context, then scaled these relative rates so that the average rate of all possible nucleotide point mutations per gene is equal to the mutation rate estimated using `MutSigCV`. We also estimated the mutation rate of oncogenic mutations in other RAS family genes and genes downstream of *KRAS*, and compared these rates to the rates measured within *KRAS*.

Selection intensities for particular amino acid substitutions were estimated by comparing the prevalence of somatic variant substitutions to the prevalence expected to occur by mutation and fixation in the absence of selection. We defined substitutions as somatic single nucleotide variants that are observed in tumors from more than one patient (recurrent) within our dataset. Quantifying mutation as occurring at an intrinsic rate *μ* per cell over the duration of tumorigenesis, the expected number of substitutions for a given site in a tumor is the product of their origination rate, *Nμ*, times the probability that the mutated lineage spreads to fixation within the tumor cell population *N*. We define this probability of fixation as *u*(*s*), where *s* is the population genetic selection coefficient, leading to a flux λ = *N*μ × *u*(*s*) of fixations of mutations. Because the probability of fixation of a neutral mutation is 1/*N* and the rate of neutral mutation within the population is *N* × μ, the rate of fixation of neutral mutations within a population is equal to μ.

The ratio of these two fluxes 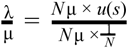 quantifies the relative importance of observed somatic variants to survival and proliferation. Therefore,

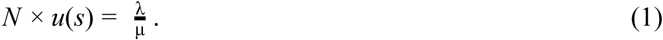

We deliberately use the term ‘selection intensity’ for the left hand side of Eq. 1, which can be estimated given knowledge of the flux of selected mutations and the intrinsic mutation rate, making a parallel to the classic derivation of ‘scaled selection coefficient’ or ‘selection intensity’ γ = 2*Ns* derived from population genetic models like the Wright-Fisher and Moran models (Sawyer and Hartl 1992; Bustamante 2005; Innan and Kim 2004; Parsons and Quince 2007). In these models the probability of fixation of a beneficial mutation is approximated to be 2*s*. Under any model of selection, the probability of fixation *u*(*s*) is a monotonically increasing function with respect to the selection coefficient of the mutant lineage. The probability of fixation *u*(*s*) can be specified in more detail by parameterization of a changing population size *N* within the extended Moran process (Moran 1958), in which the population size is not fixed (Parsons and Quince 2007). Eq. 1 supplies the selection intensity on a mutation arising in a population at a particular population size, here assumed to be consistent across tumors. More complex models of mutant allele fixation within asexually evolving populations exist—such as models that take into account multiple beneficial mutations competing for fixation (Gerrish and Lenski 1998) and multiple beneficial mutations accumulating within a single lineages competing for fixation (Desai, Fisher, and Murray 2007). However, these models require additional assumptions and additional parameters, such as the distribution of mutational effect sizes and the rate of mutations that are beneficial to cellular fitness. These two parameters are unknown for somatic tissue (Cannataro, McKinley, and St. Mary 2016).

To quantify the flux of selected mutations, we assumed that selected substitutions occur at a specific site just once during cancer evolution within a single tumor lineage, barring therapeutic intervention. This assumption is justified by 1) the extremely low per-site divergence of tumors from normal tissue (Lawrence et al. 2014), and 2) the kind of selection acting in tumorigenesis and cancer development, which is arguably acting on specific gains or losses of function (Hanahan and Weinberg 2011) rather than selecting for diversification or coevolution (Lloyd et al. 2016). Under our assumption, selection applicable to a site no longer applies after it has changed state. In this case, the actual selected flux of mutations *during selection* monotonically increases with—but is higher than would be indicated by—a Poisson calculation based solely on the number of observed variants (for an extreme example, suppose every tumor in a sample featured a somatic variant at a site: under a simple Poisson flux, this tally of fixations would yield a bounded estimate of the selected flux. However, no matter how strong the selection, the total number of observed variants would be no higher: the estimated flux should have no upper bound). Therefore, to accurately estimate selection intensity, we maximize the likelihoods of the λ*_ij_*—the Poisson rates of occurrence of fixation of specific mutation *i* at site *j* per tumor such that any tumor observed without mutation *i* at site *j* corresponds to zero substitutions, and any tumor with the somatic variant corresponds to *at least* one substitution event. We identify the value of λ*_ij_* that maximizes 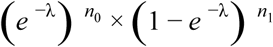, where *n*_0_ is the number of tumors without any mutation in the gene containing mutation *i* at site *j*, and *n*_1_ is the number of tumors with one substitution event *i* at site *j*.

## Results

### Deep and narrow sequencing reveals no heterogeneity within KRAS or downstream genes genes

Deep sequencing (at an average coverage of 48,163×) was performed on 27 lung tumors with detectable substitutions at the G12 codon site in KRAS and revealed no heterogeneity within each tumor among KRAS mutations (Table 2). Two of the 27 samples exhibited low-frequency variant mutations in non-*KRAS* driver genes within the RAS pathway. These low-frequency variants were a silent mutation in sample 210415AFS2, and a silent mutation and a mutation with no reported oncogenic properties (Prior, Lewis, and Mattos 2012) in sample 1071238B.

**Table 2:** Variant alleles detected with deep sequencing, sorted by G12 variant frequency.

| Sample ID | Diagnosis | Detected KRAS mutation | KRAS codons 7–15 depth of coverage (×), per run | %Variant | Other KRAS coding mutations | Other RAS-related<sup>a</sup> mutations |
| --- | --- | --- | --- | --- | --- | --- |
| S9-33 Lung | Healthy | None | NA | NA | No | No |
| S4-36 Lung | Healthy | None | NA | NA | No | No |
| S3-35 Lung | Healthy | None | NA | NA | No | No |
| 1173215F | NSCLC<sup>b</sup> | **G12C** | 251359 | 63.6 | No | No |
| 1071238B | NSCLC | G12D | 50805 | 52.6 | No | Yes<sup>c</sup> |
| 1180028F | NSCLC | **G12C** | 46484 | 52.2 | No | No |
| 1178772F | NSCLC | **G12C** | 23922 | 42.8 | No | No |
| 1180019F | LUAD<sup>d</sup> | **G12C** | 58989 | 37.9 | No | No |
| 1080556F | NSCLC | G12V | 40038 | 37.4 | No | No |
| 1179410F | NSCLC | **G12C** | 40076 | 32.3 | No | No |
| 1167759F | NSCLC | G12D | 31677 | 31.1 | No | No |
| 1091683F | NSCLC | G12D | 41192 | 30.6 | No | No |
| 1119640F | NSCLC | **G12C** | 29978 | 27.5 | No | No |
| 1170935F | NSCLC | **G12C** | 36269 | 23.3 | No | No |
| 1071289B | NSCLC | G12A | 25253 | 21.8 | No | No |
| 123877A3 | NSCLC | G12A | 24808 | 21.6 | No | No |
| 1080785F | NSCLC | G12V | 41775 | 18.4 | No | No |
| 259726A1 | NSCLC | **G12C** | 44491 | 17.6 | No | No |
| 1179420F | NSCLC | **G12C** | 44988 | 16.9 | No | No |
| 1173338F | NSCLC | **G12C** | 43231 | 16.4 | No | No |
| 1177688F | NSCLC | **G12C** | 44521 | 14.6 | No | No |
| 1119824F | NSCLC | G12D | 38212 | 12 | No | No |
| 229331A2 | NSCLC | G12D | 53742 | 11.5 | No | No |
| 1181428F | NSCLC | **G12C** | 44981 | 8 | No | No |
| 229857C2 | NSCLC | **G12C** | 43003 | 6.1 | No | No |
| 1127654F | NSCLC | G12A | 38927 | 4.9 | No | No |
| 210415AFS2 | NSCLC | G12V | 47613 | 3.5 | No | Yes<sup>e</sup> |
| 1173443F | NSCLC | **G12C** | 45364 | 2.9 | No | No |
| 1135166F | NSCLC | **G12C** | 33942 | 1.8 | No | No |
| 1119718F | NSCLC | **G12C** | 34756 | 1.7 | No | No |

<sup>a</sup>Genes sequenced at the amino acid positions reported in Table 1.<sup>b</sup>Non-Small Cell Lung Cancer<sup>c</sup>HRAS G75E at 2.3% variant frequency and HRAS D69D at 2.1% variant frequency<sup>d</sup>Lung Adenocarcinoma<sup>e</sup>BRAF L613L at 4.5% variant frequency

### Sequencing of serially pasaged xenografts reveals no heterogeneity within KRAS for G12C for mutated tumors

In addition to assessing *KRAS* heterogeneity in 27 tumors at a single time point, we tracked KRAS heterogeneity in two human lung tumors expanded in a mouse at seven distinct time points. These two patient derived xenografts were initially composed of a KRAS G12C NSCLC tumor and the surrounding human stroma included in the resected mass. As expected, when implanted in the flank of mouse, the tumor cells propagated, while the human stromal cells were rapidly replaced with mouse stromal cells. Between passages 2 and 8, which represent approximately 30 doublings of the xenograft, allele-specific PCR (Table 3) of a portion of the resected tumor (and the contaminating mouse stroma) revealed no detectable KRAS mutation other than G12C. The precise fraction of resected tumor mass containing the G12C mutation varied, most likely due to varying amounts of stromal contamination in the resected tumor.

**Table 3.** KRAS mutations, determined by allele specific PCR, in two human tumors serially passaged in mice

| Tumor | Passage | G12D | G12V | G12C | G12A | G12R | G12S | G13D | Q61R |
| --- | --- | --- | --- | --- | --- | --- | --- | --- | --- |
| Tumor 1 | 2 | <0.1% | <0.1% | 45.20% | <0.1% | <0.1% | <0.1% | <0.1% | <0.1% |
|  | 3 | <0.1% | <0.1% | >50% | <0.1% | <0.1% | <0.1% | <0.1% | <0.1% |
|  | 4 | <0.1% | <0.1% | >50% | <0.1% | <0.1% | <0.1% | <0.1% | <0.1% |
|  | 5 | <0.1% | <0.1% | >50% | <0.1% | <0.1% | <0.1% | <0.1% | <0.1% |
|  | 6 | <0.1% | <0.1% | >50% | <0.1% | <0.1% | <0.1% | <0.1% | <0.1% |
|  | 7 | <0.1% | <0.1% | >50% | <0.1% | <0.1% | <0.1% | <0.1% | <0.1% |
|  | 8 | <0.1% | <0.1% | 47.30% | <0.1% | <0.1% | <0.1% | <0.1% | <0.1% |
| Tumor 2 | 2 | <0.1% | <0.1% | >50% | <0.1% | <0.1% | <0.1% | <0.1% | <0.1% |
|  | 3 | <0.1% | <0.1% | >50% | <0.1% | <0.1% | <0.1% | <0.1% | <0.1% |
|  | 4 | <0.1% | <0.1% | >50% | <0.1% | <0.1% | <0.1% | <0.1% | <0.1% |
|  | 5 | <0.1% | <0.1% | >50% | <0.1% | <0.1% | <0.1% | <0.1% | <0.1% |
|  | 6 | <0.1% | <0.1% | >50% | <0.1% | <0.1% | <0.1% | <0.1% | <0.1% |
|  | 7 | <0.1% | <0.1% | 44.80% | <0.1% | <0.1% | <0.1% | <0.1% | <0.1% |
|  | 8 | <0.1% | <0.1% | >50% | <0.1% | <0.1% | <0.1% | <0.1% | <0.1% |

### Shallow and broad sequence analysis reveals that heterogeneity is unlikely at the time of the time treatment

Analysis of whole-exome tumor-normal sequencing of 650 LUAD tumors, 235 PAAD tumors, and 489 COADREAD tumors revealed only one COADREAD tumor (tumor sample barcode: TCGA-DY-A1H8-01A) with another KRAS mutation in addition to the G12C mutation: a silent mutation at amino acid position 20. Within the RAS-related genes and amino acid sequences (Table 1), one LUAD tumor (TCGA-05-4249-01A) contained a PIK3CA E545K mutation, one COADREAD tumor (TCGA-DM-A1D4-01A, E611E) contained a silent BRAF mutation, and four COADREAD tumors (TCGA-CM-4747-01A, E542A; TCGA-AA-A03J-01A, E545K; TCGA-AA-3696-01A, H1047R; and TCGA-DC-6683-01A, G1049S) contained PIK3CA mutations.

### Potential pathways to resistance are about as likely to arise within and downstream of KRAS. of KRAS

Deep sequencing revealed neither additional KRAS position 12 or position 61 mutations, nor the presence of oncogenic mutations upstream or downstream of KRAS within tumor samples containing KRAS position 12 mutations. Therefore, we estimated the relative rate that these mutations might arise *de novo* during treatment. First, we calculated the trinucleotide mutation profile among the lung adenocarcinoma tumors within our dataset. Applying the R package `deconstructSigs` (v1.8.0) to the observed non-recurrent single nucleotide variant mutation counts in each tumor sample that exhibited over 50 reported single nucleotide variants (excluding tumors that exhibited less than 50 reported single nucleotide variants, c.f. Rosenthal et al. 2016) provided quantification of the contribution of diverse known trinucleotide mutation signatures toward the relative trinucleotide-specific mutation rate. We averaged the distribution of trinucleotide-specific mutations profiles among all analyzed samples—generated by `deconstructSigs` using the sample-specific weight of the 30 current COSMIC signatures (http://cancer.sanger.ac.uk/cosmic/signatures)—to obtain the relative rate each of the 96 possible point mutations occur in their trinucleotide context. Substitutions were characterized by a relatively high frequency of G→T mutations, a mutational signature common in lung adenocarcinomas (Fig. 2; shown here in the equivalent C→A context), likely caused by mutagens in tobacco (Porta et al. 2009).

**Figure 2:**
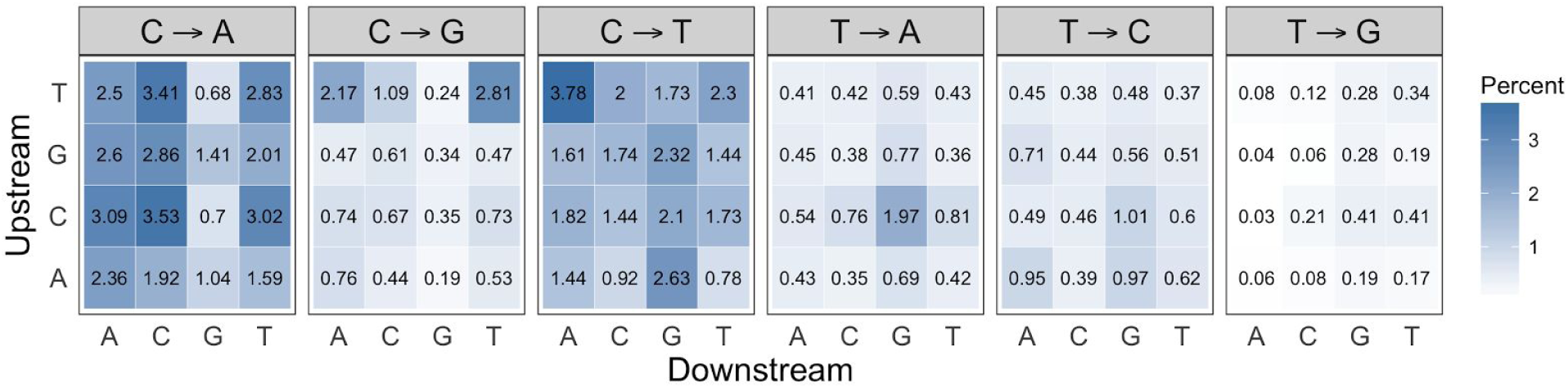
Heatmap of the average trinucleotide mutation profile within our lung adenocarcinoma dataset. Darker blue corresponds to higher mutation rates, and the percent of mutations represented by each of the 96 trinucleotide contexts are displayed within the tiles.

Next, we used MutSigCV to estimate the average site-specific nucleotide mutation rate within *KRAS* in our dataset. We found that the average rate that mutations occurred in *KRAS* from tumorigenesis to resection was 4.7 × 10^-6^ per nucleotide. We calculated the rate at which mutations arise at the KRAS G12C site by distributing this average *KRAS* nucleotide site mutation rate among each observed nucleotide transition and transversion (Fig. 3). Using these rates, we then calculated the rate of specific amino acid mutations at this site given the distribution of specific nucleotide mutations (Fig. 4). We found that mutations in phenylalanine are expected to occur at the highest rate, followed by serine, tyrosine, mutation to a stop codon, cysteine, arginine, tryptophan, and glycine, respectively. Although mutations at KRAS position 12 to phenylalanine, tyrosine, and tryptophan are not observed clinically (they cannot be produced by single nucleotide mutations from the wild type GGT codon), it is likely that they will be oncogenic: *in vitro* experiments (Seeburg et al. 1984) and analyses of KRAS protein structure (Scheffzek et al. 1997) suggest that any other substitution at the position 12 site—other than proline—would result in the constitutive activation associated with tumorigenesis. Mutations to a stop codon, cysteine, or glycine will not rescue the tumor phenotype in the context of an effective KRAS G12C therapeutic, as they result in a nonsense mutation, a silent mutation, and reversion to the amino acid encoded by the human reference sequence, respectively. At amino acid position 61 our entire dataset contained only two recurrent substitutions: three instances each of Q61L and Q61H, occurring at estimated mutation rates of 9.5 × 10^−7^ and 1.9 × 10^−6^, respectively.

**Figure 3:**
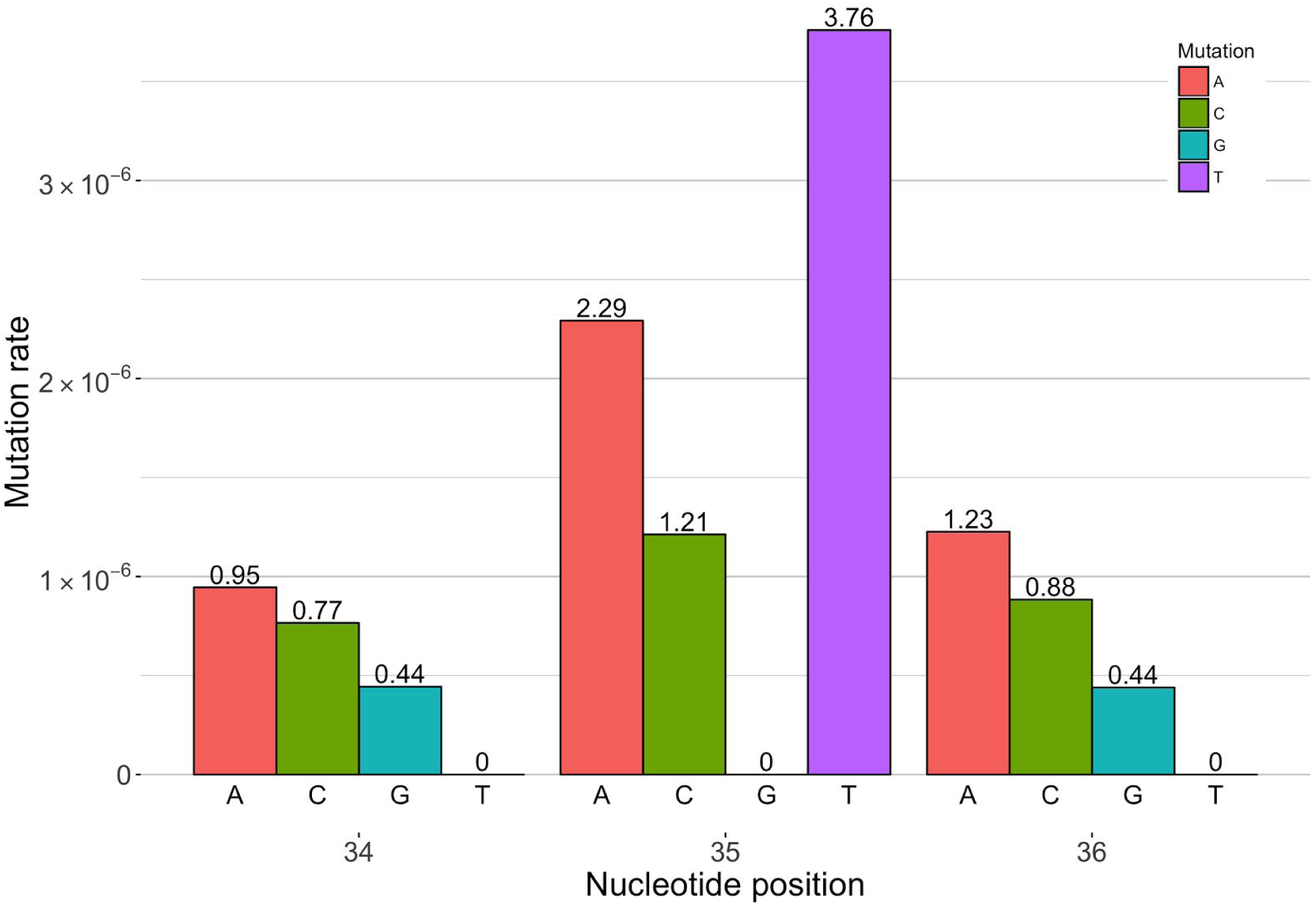
The nucleotide point mutation rate from tumorigenesis to resection at *KRAS* nucleotide positions 34, 35, and 36, calculated for those sites with the position 34 G→T mutation, responsible for the G12C amino acid replacement, present.

**Figure 4:**
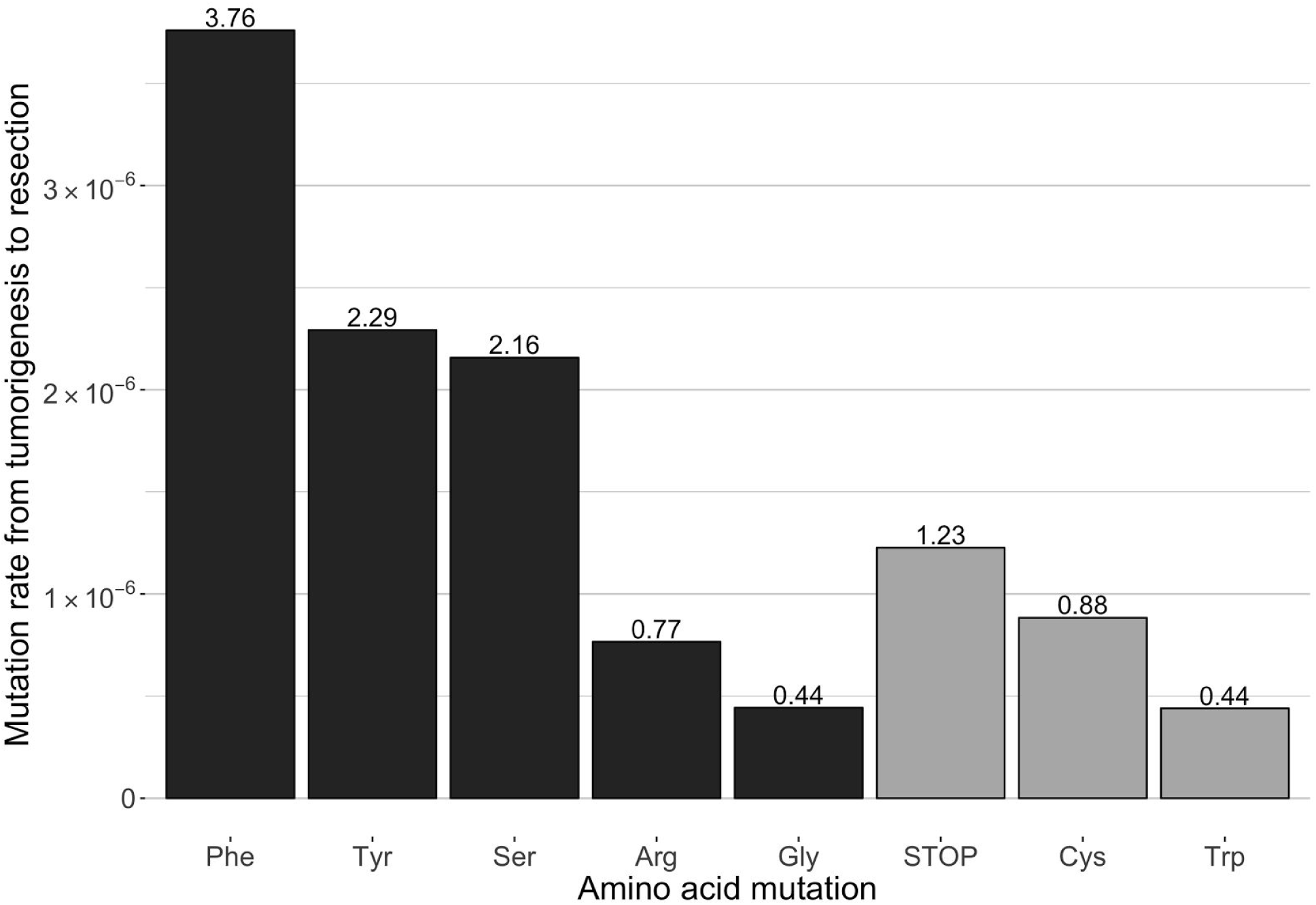
The mutation rates of G12C mutant KRAS at amino acid position 12. The net rate of mutation is 1.2 × 10^-5^ per nucleotide from tumorigenesis to resection. Mutations encoding cysteine would remain targeted by the therapeutic. Mutations encoding a premature STOP codon would create a nonfunctional and non-oncogenic protein. Mutations encoding a glycine amino acid would result in reversion to the human reference amino acid at position 12 of KRAS.

Next, we calculated the rate of mutation within the other *RAS* family genes, *NRAS* and *HRAS*, as well as genes downstream of *KRAS* that may rescue the tumor phenotype, namely *PIK3CA, MAP2K1, MAP2K2, MAPK1, MAPK3, AKT1*, and *MTOR (Huang and Fu 2015; Roberts and Stinchcombe 2013)*, using the same methodology described for the within-KRAS analysis. We are only concerned with mutations that might rescue the tumor phenotype, and thus only calculated the mutation rate of recurrently observed point mutations within our dataset (Fig. 5A). We found that the overall rate of oncogenic mutations in related RAS genes and downstream genes (1.2 × 10^−5^) was about the same as the mutation rate of oncogenic mutations (Tyr, Phe, Ser, Arg, Trp) at KRAS position 12 (9.4 × 10^-6^) and KRAS position 61 (2.9 × 10^−6^) combined (1.2 × 10^−5^). The rate of mutation leading to a particular recurrent substitution within the downstream genes does not exhibit a statistically significant correlation with the frequency that the substitution is observed (*r* = 0.14, *P* = 0.64). This lack of correlation is presumably a consequence of subsequent selection on nucleotide state. Therefore, we estimated the selective pressure on each observed substitution.

**Figure 5:**
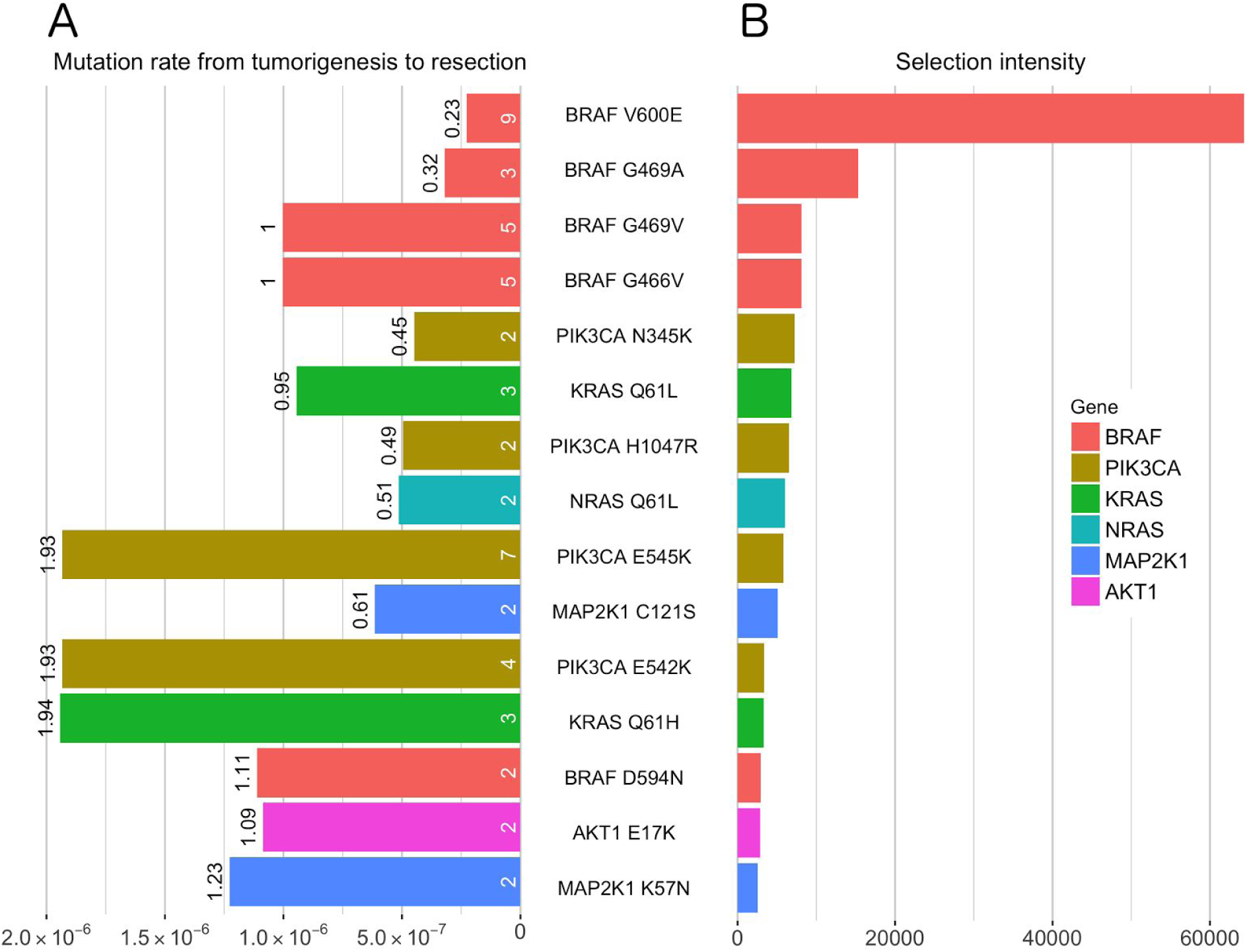
The mutation rate, frequency of, and estimated selection intensity for downstream variants that may result in resistance to targeted KRAS G12C therapy. (A) The mutation rate at mutant KRAS position Q61, and at sites in genes downstream of RAS that also exhibit recurrent mutations among lung adenocarcinomas in the TCGA and Yale-Gilead datasets. The numbers within the bars indicate the number of mutations observed in 650 exome sequence lung adenocarcinoma primary tumors. (B) The estimated selection intensities for KRAS Q61 mutations and for mutations in genes downstream of RAS that also exhibit recurrent mutations among all sequenced lung adenocarcinomas in the TCGA and Yale-Gilead datasets.

### Estimates of selection intensity informed predictions of likely pathways to resistance

We calculated the selection intensity (Eq. 1) for recurrent KRAS Q61 mutations and recurrent substitutions in associated genes (Fig. 5B). The BRAF V600E mutation yielded the highest selection intensity, suggesting that it is the mutation most likely to confer the largest fitness advantage to cancer cells.

## Discussion

Here we have conducted deep sequencing of multiple NSCLC tumors, demonstrating that oncogenic mutations that might compromise targeted treatment of KRAS G12C positive tumors, both at the KRAS G12 site, the KRAS Q61 site, and among related genes in the RAS pathway, are unlikely to be present at the time of treatment. Additionally, we found that eight experimental passages of two tumor lines as PDXs in mice, constituting greater than 30 tumor doublings each, introduced no detectable heterogeneity within *KRAS*. Moreover, we estimated the rates of *de novo* mutations arising after the initiation of treatment, revealing a similar rate of mutation within genes downstream of KRAS when compared to oncogenic mutations at KRAS positions 12 and 61. Finally, we estimated the selection intensity on all of these oncogenic mutations, conveying a quantitative measure of their importance to tumorigenesis and cancer development.

Our results indicate that targeted KRAS G12C therapy shows potential for durable effectiveness given the low likelihood of extant mechanisms of drug resistance. Synthetic lethal therapy for oncogenic *KRAS* mutants also show promise (Mao et al. 2014), and inhibitory drugs targeting proteins encoded by *PIK3CA, AKT1*, *BRAF*, and other genes within the RAS pathway, are either currently available or in phase I–III trials (Huang and Fu 2015), indicating that many likely pathways to resistance of targeted KRAS G12C therapy are potentially treatable by serial or simultaneous therapeutics. Of all potential mutations, the BRAF V600E and G469A mutations exhibited the highest selection intensities. Thus, they would be predicted to confer the largest fitness advantage to resistant subclones. Remarkably, these two mutations that we estimated to have the highest cancer effect sizes for NSCLC were the only two mutations that were found to confer resistance to targeted therapy of *EGFR*, a gene directly upstream of *KRAS*, in lung tumor cell lines (Ohashi et al. 2012).

Our estimates of mutation rates and selection intensities inform clinicians about the rates that different resistance mechanisms may arise, and the relative effect those mechanisms will have on tumor relapse. Although analyses described here have been applied to specifically evaluate the selective dynamics of mutations within and downstream of KRAS, these analyses can be performed on any tissue and genetic pathway with extensive tumor sequence data. Furthermore, as new methodologies are developed and increased data is collected that elucidates the complexities of in somatic evolution in each person—such as those imposed by germ-line polymorphisms (Zienolddiny et al. 2005) or those that arise naturally through ontogenic changes to the tumor microenvironment (Rozhok and DeGregori 2016)—it should be possible to develop additional theory to incorporate them by modulating intrinsic rates of mutation or probabilities of fixation. We have provided an evolutionary framework with potential utility in prediction of pathways to resistance as new targeted therapies become available.

One relevant complexity that we assumed is constancy of selective effect across the duration of tumorigenesis, cancer development, and medical intervention. We assumed that the profile of mutations and the selection intensity on specific mutations during KRAS G12C treatment is the same as the selection intensity estimated from the period of progression of tumors from healthy to cancerous tissue in our lung adenocarcinoma dataset. However, it is generally understood that a tumor under treatment may present a different selective landscape than an untreated tumor, and a different landscape entails different selection for yet-unknown variants. For instance, the EGFR T790M mutation is a known oncogenic agent in lung adenocarcinoma, but is rarely detected in untreated tumors. In contrast, the T790M variant is responsible for approximately 50% of acquired resistance to EGFR tyrosine kinase inhibitors in EGFR L858R positive tumors because the T790M variant increases ATP affinity of EGFR L858R mutants by more than an order of magnitude, and the L790M/L858R double mutants also show more oncogenic phosphorylation activity when compared to the L858R or T790M mutants alone (Yun et al. 2008; Suda et al. 2009; Mulloy et al. 2007). Thus, the selective pressure for the EGFR T790M mutation varies depending on the mutational background and treatment regime of the tumor in which it arises. Similarly, we may not be aware of novel KRAS mutations that are selected in the presence of yet-untested KRAS-specific therapies. Resistance to EGFR therapy is also conferred by other EGFR mutations and mutations downstream of EGFR, and thus our analysis points to important potential mechanisms of *KRAS* specific therapy using the data currently available.

Our analysis constitutes the first estimate of the intensity of selection on point mutations within somatic tissue from whole exome sequencing data. The estimates of selection intensity provided by this evolutionary framework are a function of the relative fitness advantage conferred by the mutations, and thus offer insight into the degree that specific mutations drive the growth and evolution of cancer, and into the therapeutic potential of targeted therapeutics that are able to fully abrogate their aberrant function.

## Author contributions

VLC and JPT developed new methods and approaches. VLC and SGG performed all computational analyses. AEG performed experiments. VLC, SGG, CS, ZZ, MP, AEG, and JPT wrote the manuscript.

## Data and materials availability

Data collected and scripts used in this analysis will be made available upon publication. Data retrieved from the Broad Institute are available from Broad Institute TCGA Genome Data Analysis Center (2016): Firehose stddata__2016_01_26 run. (Broad Institute 2016)

